# Meta-analytic activation maps can help identify affective processes captured by contrast-based task fMRI: the case of threat-related facial expressions

**DOI:** 10.1101/820969

**Authors:** M. Justin Kim, Annchen R. Knodt, Ahmad R. Hariri

## Abstract

Meta-analysis of functional magnetic resonance imaging (fMRI) data is an effective method for capturing the distributed patterns of brain activity supporting discrete cognitive and affective processes. One opportunity presented by the resulting meta-analysis maps (MAMs) is as a reference for better understanding the nature of individual contrast maps (ICMs) derived from specific task fMRI data. Here, we compared MAMs from 148 neuroimaging studies representing the broad emotion categories of fear, anger, disgust, happiness, and sadness with ICMs from fearful > neutral and angry > neutral facial expressions from an independent dataset of task fMRI (*n* = 1263). Analyses revealed that both fear and anger ICMs exhibited the greatest pattern similarity to fear MAMs. As the number of voxels included for the computation of pattern similarity became more selective, the specificity of MAM-ICM correspondence decreased. Notably, amygdala activity long considered critical for processing threat-related facial expressions was neither sufficient nor necessary for detecting MAM-ICM pattern similarity effects. Our analyses suggest that both fearful and angry facial expressions are best captured by distributed patterns of brain activity associated with fear. More generally, our analyses demonstrate how MAMs can be leveraged to better understand affective processes captured by ICMs in task fMRI data.

## Introductions

Understanding how emotions map onto human brain function is a long-standing aim of affective neuroscience. To achieve this goal, affective neuroscientists have heavily employed functional magnetic resonance imaging (fMRI) to examine whether different properties of emotion based on existing theories – such as valence/arousal dimension (Russell, 1980) or basic emotion categories (Ekman, 1992) – may be reflected in patterns of brain activity. Early fMRI studies that aimed to elucidate the neural representations of categorical emotions focused on the amygdala because of its demonstrated importance in aversive learning as exemplified through the acquisition of a conditioned fear response (LeDoux, 1993; Maren, 2001). However, fMRI studies yielded mixed results wherein amygdala activity was not only elicited by threat-related emotions (e.g., fear, anger) but also other categories of emotion (e.g., happiness, sadness) (Davis & Whalen, 2001).

In fact, it has been suggested that amygdala activity alone does not provide a sufficient level of specificity in distinguishing emotion categories (Fitzgerald et al., 2006). More generally, fMRI studies examining discrete emotion categories have not revealed correspondingly discrete brain regions (Lindquist et al., 2012). In contrast, recent research employing multivoxel pattern analysis (MVPA) offers evidence that discrete emotion categories may be best represented by distributed patterns of activity across the brain (Kassam et al., 2013; Kragel & Labar, 2015; Kragel et al., 2019; Peelen et al., 2010; Saarimäki et al., 2016; but see Barrett & Satpute, 2019).

A rapidly expanding portfolio of fMRI studies has allowed for a series of computational methods designed to generate voxel-wise meta-analysis maps (MAMs) of brain activity associated with specific cognitive and affective processes (Eickoff et al., 2009; Kober & Wager, 2010; Yarkoni et al., 2011). MAMs typically share the same coordinate or stereotaxic space (e.g., MNI or Talairach) as individual contrast maps (ICMs), which represent subject-level brain activity associated with a study-specific contrast of interest. As such, MAMs could provide a reference map for a given mental process or behavior. For example, ICMs of a typical emotion category, such as fear, should show patterns of brain activity more similar to MAMs of the corresponding category (i.e., fear) than other categories (e.g., disgust or sadness). In other words, if affective information for fear is indeed represented in brain activity for fear ICMs, they should correspond to the associated MAMs for fear. This not only presents a testable prediction but also an opportunity to refine ICMs (through pattern similarity analysis with MAMs) in a fashion that maximizes their ability to capture brain activity supporting a specific mental process.

In this study, we aimed to test a simple idea: would an ICM of a given emotion category show greater similarity to the MAM of the same category over others? There are at least three conditions that serves as a prerequisite for such an examination: 1) MAMs from voxel-wise meta-analyses of fMRI data, 2) category-specific ICMs from a study with a sufficiently large sample size, and 3) independence between the MAMs and ICMs – that is, the ICMs selected for testing should not have been used in the generation of the MAMs. Here, we take advantage of datasets that satisfy these conditions. First, Wager and colleagues (2015) have conducted a computational meta-analysis of 148 functional neuroimaging studies to generate MAMs for five emotion categories: fear, anger, disgust, happiness, and sadness. Second, the Duke Neurogenetics Study (DNS) offers a large independent dataset (*n* = 1263) of ICMs for fear, anger, surprise, and neutral emotions from a widely-utilized face-matching task. Importantly, none of the ICMs from the DNS were included in the generation of MAMs.

Based on the existing literature, we hypothesized that ICMs for fear and anger would show the greatest pattern similarity to MAMs for fear and anger, respectively. Moreover, we sought to examine the possibility that affective information pertaining to emotion categories is distributed across the whole brain by systematically varying the number of voxels submitted for pattern similarity analysis. Finally, given the prominence of the amygdala in the affective neuroscience literature (Adolphs et al., 1995; Costafreda et al., 2008; Kim et al., 2011; Phelps & LeDoux, 2005), we tested whether amygdala activity specifically was either sufficient or necessary to produce the MAM-ICM pattern similarity observed in whole brain analyses.

## Methods

### Participants

1263 participants (717 women, 19.7 ± 1.3 years of age) successfully completed the Duke Neurogenetics Study (DNS) between January 2010 and November 2016 including an fMRI task eliciting threat-related brain activity. All participants provided written informed consent according to the Duke University Medical Center Institutional Review Board. To be eligible for the DNS, participants were required to be free of the following conditions: 1) medical diagnoses of cancer, stroke, head injury with loss of consciousness, untreated migraine headaches, diabetes requiring insulin treatment, chronic kidney, or liver disease; 2) use of psychotropic, glucocorticoid, or hypolipidemic medication; and 3) conditions affecting cerebral blood flow and metabolism (e.g., hypertension). As DNS followed a standardized procedure, we note that the following description of the methods is also described elsewhere (e.g., Kim et al., 2018).

### Face Matching Task

The face matching task used in the DNS consisted of four task blocks interleaved with five control blocks. A total of four emotion categories were used for each task block: fear (F), anger (A), surprise (S), and neutral (N), taken from a standardized facial expression set (Ekman & Friesen, 1976). Participants viewed the task blocks in one of four randomly assigned orders as determined by a Latin Square (i.e., FNAS, NFSA, ASFN, SANF). During task blocks, participants viewed a trio of faces and matched one of two faces identical to a target face. Each trial in the task blocks lasted for 4 s with a variable interstimulus interval of 2-6 s (mean = 4 s), for a total block length of 48 s. The control blocks consisted of six geometric shape trios, which were presented for 4 s with a fixed interstimulus interval of 2 s for a total block length of 36 s. Each block was preceded by a brief instruction (“Match faces” or “Match shapes”) lasting 2 s. Total task time was 390 s.

### fMRI Data Acquisition

Each participant was scanned using one of the two identical research-dedicated GE MR750 3T scanner equipped with high-power high-duty-cycle 50-mT/m gradients at 200 T/m/s slew rate, and an eight-channel head coil for parallel imaging at high bandwidth up to 1MHz at the Duke-UNC Brain Imaging and Analysis Center. A semi-automated high-order shimming program was used to ensure global field homogeneity. A series of 34 interleaved axial functional slices aligned with the anterior commissure-posterior commissure plane were acquired for full-brain coverage using an inverse-spiral pulse sequence to reduce susceptibility artifacts (TR/TE/flip angle=2000 ms/30 ms/60; FOV=240mm; 3.75×3.75×4mm voxels; interslice skip=0). Four initial radiofrequency excitations were performed (and discarded) to achieve steady-state equilibrium. To allow for spatial registration of each participant’s data to a standard coordinate system, high-resolution three-dimensional T1-weighted structural images were obtained in 162 axial slices using a 3D Ax FSPGR BRAVO sequence (TR/TE/flip angle = 8.148 ms / 3.22 ms / 12°; voxel size=0.9375×0.9375×1mm; FOV=240mm; interslice skip=0; total scan time = 4 min and 13 s). In addition, high-resolution structural images were acquired in 34 axial slices coplanar with the functional scans and used for spatial registration for participants without Ax FSPGR BRAVO images (TR/TE/flip angle=7.7 s/3.0 ms/12; voxel size=0.9×0.9×4mm; FOV=240mm, interslice skip=0).

### fMRI Data Preprocessing

Anatomical images for each subject were skull-stripped, intensity-normalized, and nonlinearly warped to a study-specific average template in the standard stereotactic space of the Montreal Neurological Institute (MNI) template using ANTs (Klein et al., 2009). BOLD time series for each subject were processed in AFNI (Cox, 1996). Images for each subject were despiked, slice time-corrected, realigned to the first volume in the time series to correct for head motion, coregistered to the anatomical image using FSL’s Boundary Based Registration (Greve & Fischl, 2009), spatially normalized into MNI space using the non-linear warp from the anatomical image, resampled to 2 mm isotropic voxels, and smoothed to minimize noise and residual difference in gyral anatomy with a Gaussian filter, set at 6-mm full-width at half-maximum. All transformations were concatenated so that a single interpolation was performed. Voxel-wise signal intensities were scaled to yield a time series mean of 100 for each voxel. Volumes exceeding 0.5 mm framewise displacement (FD) or 2.5 standardized temporal derivative of RMS variance over voxels (DVARS)(Nichols, 2017; Power et al., 2014) were censored.

### Individual Contrast Maps

The AFNI program *3dREMLfit* (http://afni.nimh.nih.gov/) was used to fit a general linear model for first-level fMRI data analyses. To obtain emotion-specific parameter estimates, we explicitly modeled each respective task block (convolved with the canonical hemodynamic response function) along with the adjacent half of the preceding and following control blocks, and a first order polynomial regressor to account for low frequency noise. This allowed for the estimation of the individual task block parameters while minimizing the influence of adjacent task blocks as well as low frequency noise across the entire run. The resulting parameter estimates for the fear and anger task blocks and the neutral task blocks were then subtracted to obtain the fearful > neutral and angry > neutral faces ICMs (henceforth referred to as ICM-F and ICM-A, respectively), and these ICMs were used to compute pattern similarity with the MAMs. The contrast surprise > neutral was omitted from the present study, because a corresponding surprise MAM does not exist. ICMs were then used in second-level random effects models in SPM12 (http://www.fil.ion.ucl.ac.uk/spm) accounting for scan-to-scan and participant-to-participant variability to determine mean emotion-specific activity using one-sample *t*-tests. A statistical threshold of *p* < 0.05, family-wise error (FWE)-corrected across the whole brain was applied to the fearful > neutral and angry > neutral contrasts, respectively.

### fMRI Quality Assurance Criteria

Quality control criteria for inclusion of a participant’s imaging data were: > 5 volumes for each condition of interest retained after censoring for framewise displacement (FD) and DVARS and sufficient temporal SNR within the bilateral amygdala, defined as no greater than 3 standard deviations below the mean of this value across participants. The amygdala was defined using a high-resolution template generated from the 168 Human Connectome Project datasets (Tyszka & Pauli, 2016). Additionally, data were only included in further analyses if the participant demonstrated sufficient engagement with the task, defined as achieving at least 75% accuracy during the task blocks.

### Meta-Analysis Maps

The MAMs utilized in the present study are based on a meta-analysis of 148 fMRI studies of emotional categories (Wager et al., 2015). The MAMs were generously made available by these study authors on their website (https://canlabweb.colorado.edu/fmri-resources.html). In brief, the original meta-analysis consisted of five distinct emotion categories (fear, anger, disgust, sadness, happiness), and MAMs for each category were generated in standard MNI space using a hierarchical Bayesian model that summarizes the expected frequency of activation for a given emotion category. The values that each voxel of the MAMs assumes is reflective of this information, such that taking the integral over any area of the brain represents the expected number of activation centers for all studies of a given emotion category (Wager et al., 2015). The overall findings indicated that brain activity patterns that are diagnostic of distinct categories of emotion are characterized as widespread (i.e., distributed not only across multiple brain regions, but also many neural systems that span cognitive, perceptual, and motor functions). For the purposes of the present study, higher intensity voxel values of the MAMs indicate a greater likelihood with which the MAMs correspond to a given emotion category. Importantly, none of the 148 studies that were included in this meta-analysis overlapped with the DNS, ensuring independence across MAMs and ICMs (the MAMs included studies published from 1993 to 2011; the first study using the ICMs from the emotional face matching task in the DNS was published in 2012).

### MAM-ICM Pattern Similarity Computation

For quantifying MAM-ICM pattern similarity, we adopted a modified version of an approach described by Shahane and colleagues (2019). First, prior to each analysis, ICMs were masked with the target MAMs, to match the number of non-zero voxels. Then, for each MAM-ICM pair, all non-zero voxels were vectorized and demeaned in order to compute their correlation coefficients, which were subsequently converted to *z* scores using Fisher’s *r*-to-*z* transformation (Figure 1). Computation of correlation coefficients across vectorized voxels was achieved with *3ddot* implemented within AFNI (Cox, 1996). Higher *z* scores indicated greater pattern similarity between a given MAM-ICM pair. Since there were 5 MAMs, a total of 5 *z* scores were computed for each of two ICMs for each participant. As we were primarily interested in how well a given ICM (e.g., ICM-F) aligned with its corresponding MAM (e.g., MAM-F) above and beyond other MAMs, the *z* scores were compared across the MAMs. Two models were generated to test this: 1) a one-way repeated measures analysis of variance (ANOVA) where *z* scores for a given ICM and MAMs for each of the five emotion categories were compared, and 2) a paired *t*-test where *z* scores for a given ICM and MAMs for fear and anger, specifically, were compared. The latter model was used to specifically focus on the two emotion categories that were available as both ICMs and MAMs. Analyses were done separately for ICM-F and ICM-A. In this way, we could confirm the *a priori* prediction that ICM-F would correspond better to the MAM-F, above and beyond the MAMs of other emotion categories; then, the same procedure was applied to test whether ICM-A would show increased pattern similarity to the MAM-A compared to other MAMs.

**Figure 1.**
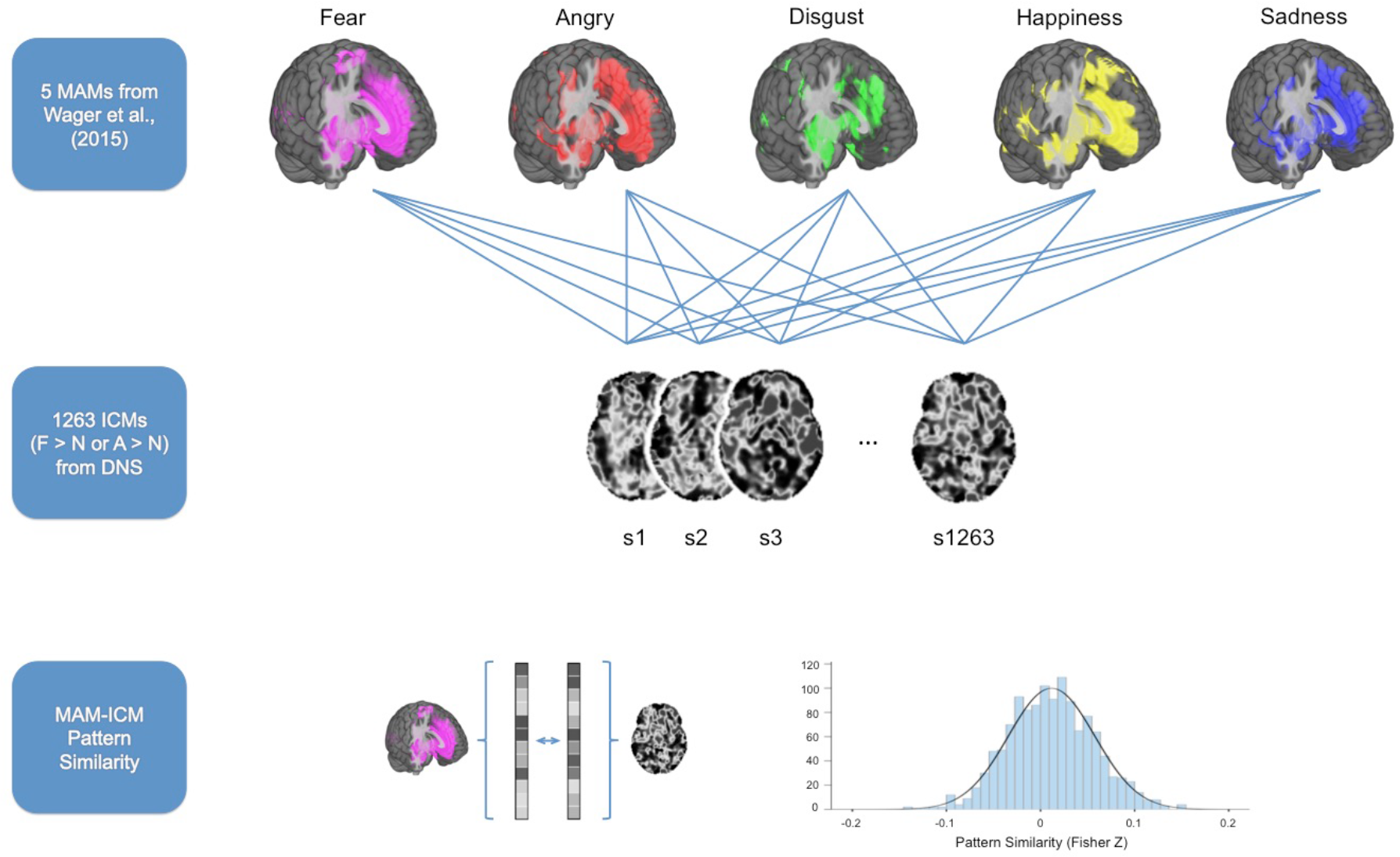
Summary of data analysis procedures used for assessing MAM-ICM pattern similarity. ICMs for fearful > neutral and angry > neutral derived from 1263 performing an emotional face matching task were compared with five MAMs corresponding to different categories of emotion from Wager et al. (2015). For each MAM-ICM pair, all non-zero voxels were vectorized in order to compute their correlation coefficient, which was subsequently converted to *z* scores using Fisher’s *r*-to-*z* transformation.

To test whether the number of voxels included in the analyses impacted the results, MAM-ICM pattern similarity measures were computed repeatedly using MAMs at different thresholds. The threshold was systematically varied, ranging from unthresholded to 0.1 (intensity values), with each step increasing the threshold by twofold (unthresholded – 0.001 – 0.005 – 0.01 – 0.05 – 0.1). It is notable that there was a general tendency for cortical areas to become reduced as a function of increased MAM threshold, while the amygdala was among the last regions to remain, regardless of emotion category (Figure 2). Effect sizes from the analyses that compared a given ICM across different MAMs (partial *η*^2^ for ANOVA, Cohen’s *d* for *t*-test) were used to describe the effect of applying different thresholds on the pattern similarity metrics.

**Figure 2.**
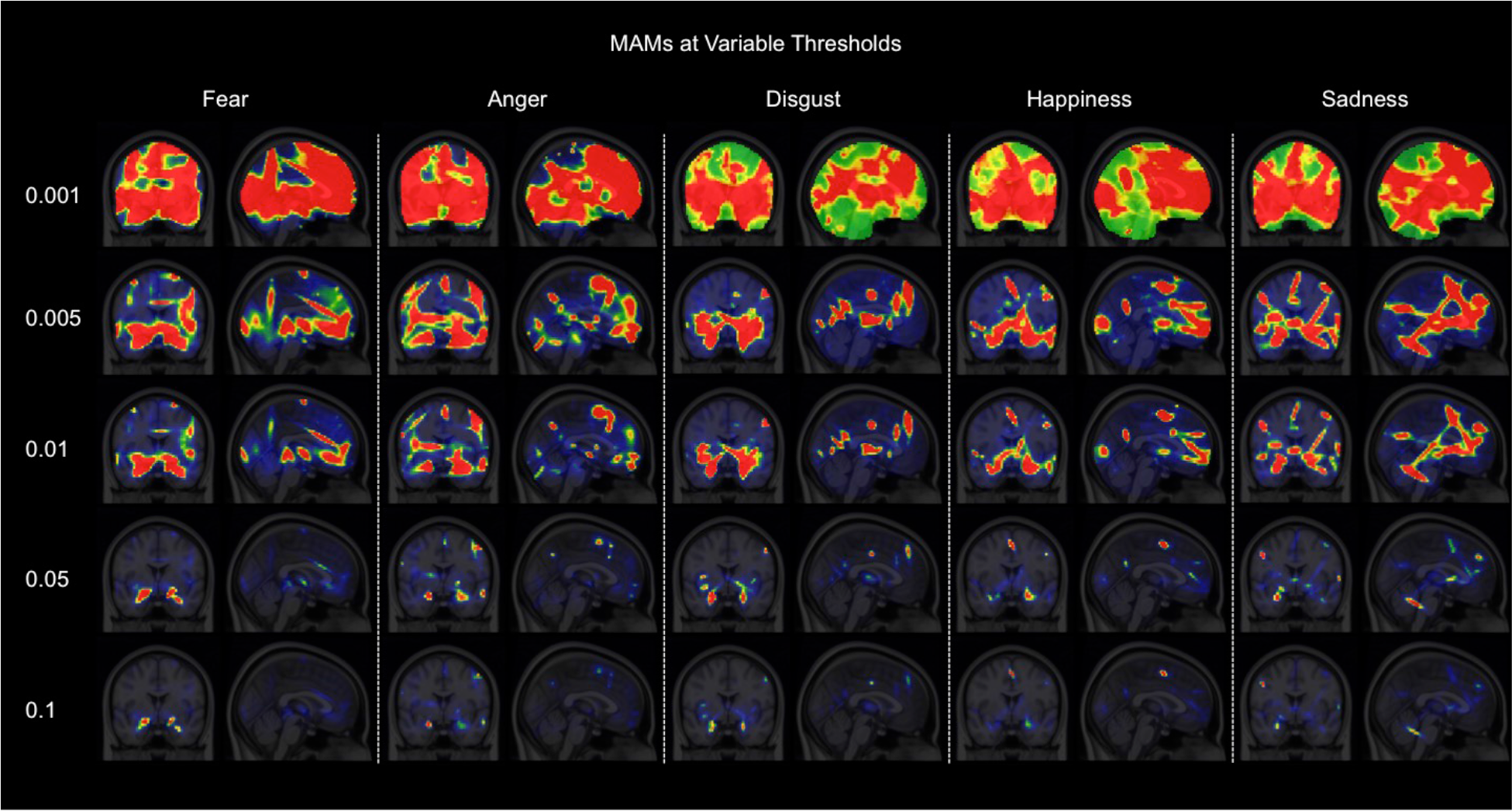
Five emotion-specific MAMs at systematically varied thresholds. Voxels that survived the increasingly stringent thresholds (i.e., the higher values noted on the left-hand side) are depicted by hot colors.

Finally, to test the contribution of amygdala activity specifically on the pattern similarity results from the main analyses, two sets of subsequent analyses were performed on the data. First, each MAM-ICM pair was masked with an anatomical ROI of the amygdala, divided into basolateral and centromedial subregions (Tyszka & Pauli, 2016), which then underwent the same procedure described above using only the amygdala voxels (here, only unthresholded MAM voxels was used). In a separate set of analyses, each MAM-ICM pair was masked with a reversed mask of the amygdala ROI, such that all amygdala voxels were removed from further analysis. Then, the same procedure as the main analyses was repeated for the amygdala-excluded MAM-ICM pairs. The latter analysis tested the prediction that if the amygdala voxels contain important information about distinct emotion categories, then it would yield decreased pattern similarity metrics for corresponding MAM-ICM pairs.

## Results

### ICMs: Fearful > Neutral and Angry > Neutral

Across the entire brain, both ICMs yielded significant activity in the amygdala, supramarginal gyrus/angular gyrus extending to the superior temporal sulcus (STS), and inferior frontal gyrus (IFG). The fearful > neutral ICMs also revealed significantly increased activity in the occipital pole and the inferior temporal gyrus (ITG). Significantly activated amygdala voxel clusters were isolated within the amygdala proper and not a part of a larger cluster that extends to other brain regions (Figure 3, Table 1).

**Figure 3.**
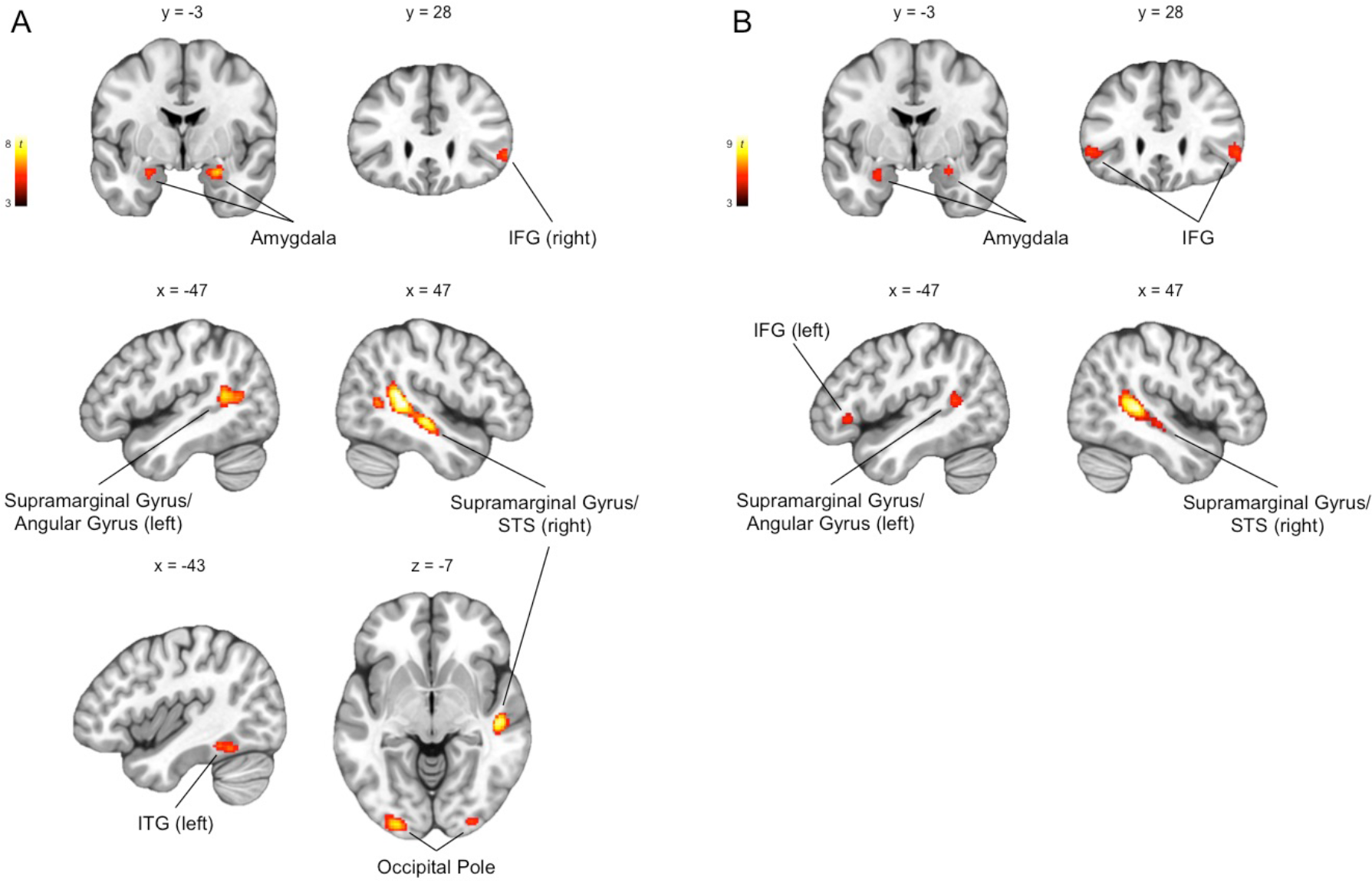
Group-level whole brain responses from 1263 participants performing an emotional face matching task (*p* < 0.05, FWE-corrected for the whole brain; *k* ≥ 30 are visualized). (A) Brain regions that showed significantly increased activity to fearful > neutral included the amygdala, supramarginal gyrus/angular gyrus/superior temporal sulcus (STS), inferior frontal gyrus (IFG), inferior temporal gyrus (ITG), and the occipital pole. (B) Similar brain regions showed significantly increased activity to anger > neutral.

**Table 1.**
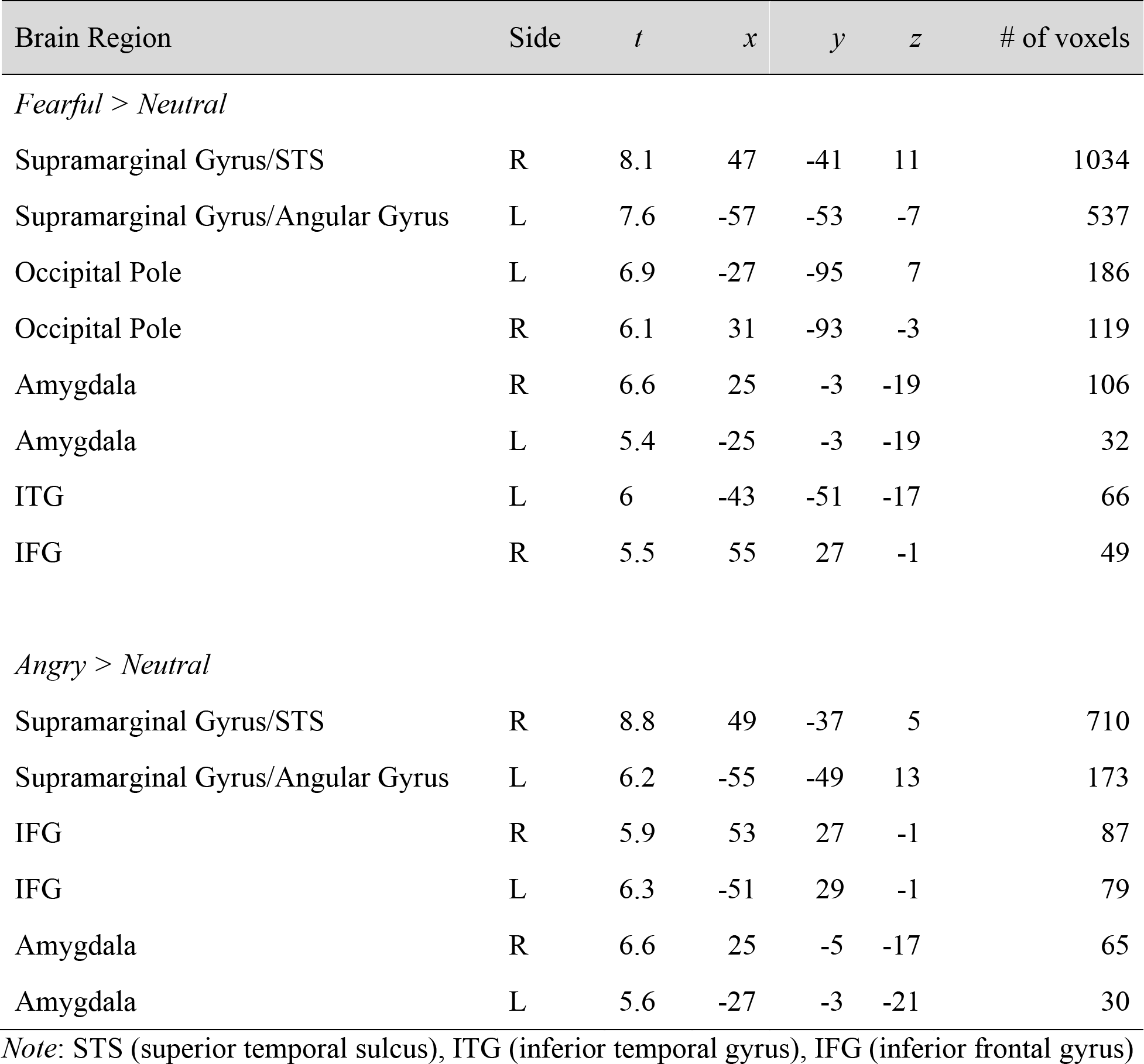
Brain regions showing significant activation for the contrasts of fearful > neutral and angry > neutral (*p* < 0.05, FWE-corrected for the whole brain).

### Whole Brain MAM-ICM Pattern Similarity for Fear

Overall, there was significant pattern similarity between ICM-F and MAM-F (*M* = 0.0065, *SD* = 0.03, [min, max] = [−0.11, 0.11] ; one-sample *t*-test: *t*_(1262)_ = 7.57, *p* < 0.000001, *d* = 0.21). This effect remained significant when the MAMs were thresholded at varying levels (all *p*s < 0.002), except for one instance (ICM-F and MAM-F pair thesholded at 0.1; *p* = 0.16).

Repeated measures ANOVA showed significant differences of MAM-ICM pattern similarity for ICM-F across the five MAMs (*F*_(4,5048)_ = 17.27, *p* < 0.000001; partial *η*^2^ = 0.014). *Post hoc* analysis revealed that pattern similarity for MAM-F was significantly greater than the other four MAMs (all *p*s < 0.002). This finding remained when the threshold was increased to 0.01 (i.e., less voxels were selected); however, this effect was no longer observable when the threshold was further increased. Paired *t*-tests showed similar findings as the ANOVA, such that pattern similarity between ICM-F and MAM-F was significantly greater than MAM-A (*t*_(1262)_ = 3.3, *p* = 0.001; *d* = 0.09). Again, this effect was observable until the threshold was increased to 0.01. When the threshold was set to the highest level (0.1), an unexpected opposite effect was found such that pattern similarity between ICM-F and MAM-F was significantly less than for ICM-F and MAM-A (*t*_(1262)_ = −2.65, *p* = 0.008; *d* = 0.1).

Effect sizes of the ANOVA and *t*-test results gradually decreased as a function of increased threshold levels. As ICM-F showed the highest level of pattern similarity to MAM-F over other MAMs, diminishing effect sizes suggest a relative decrease in the specificity of ICM-F to MAM-F. These findings are summarized in Figure 4 (white bars and circles).

**Figure 4.**
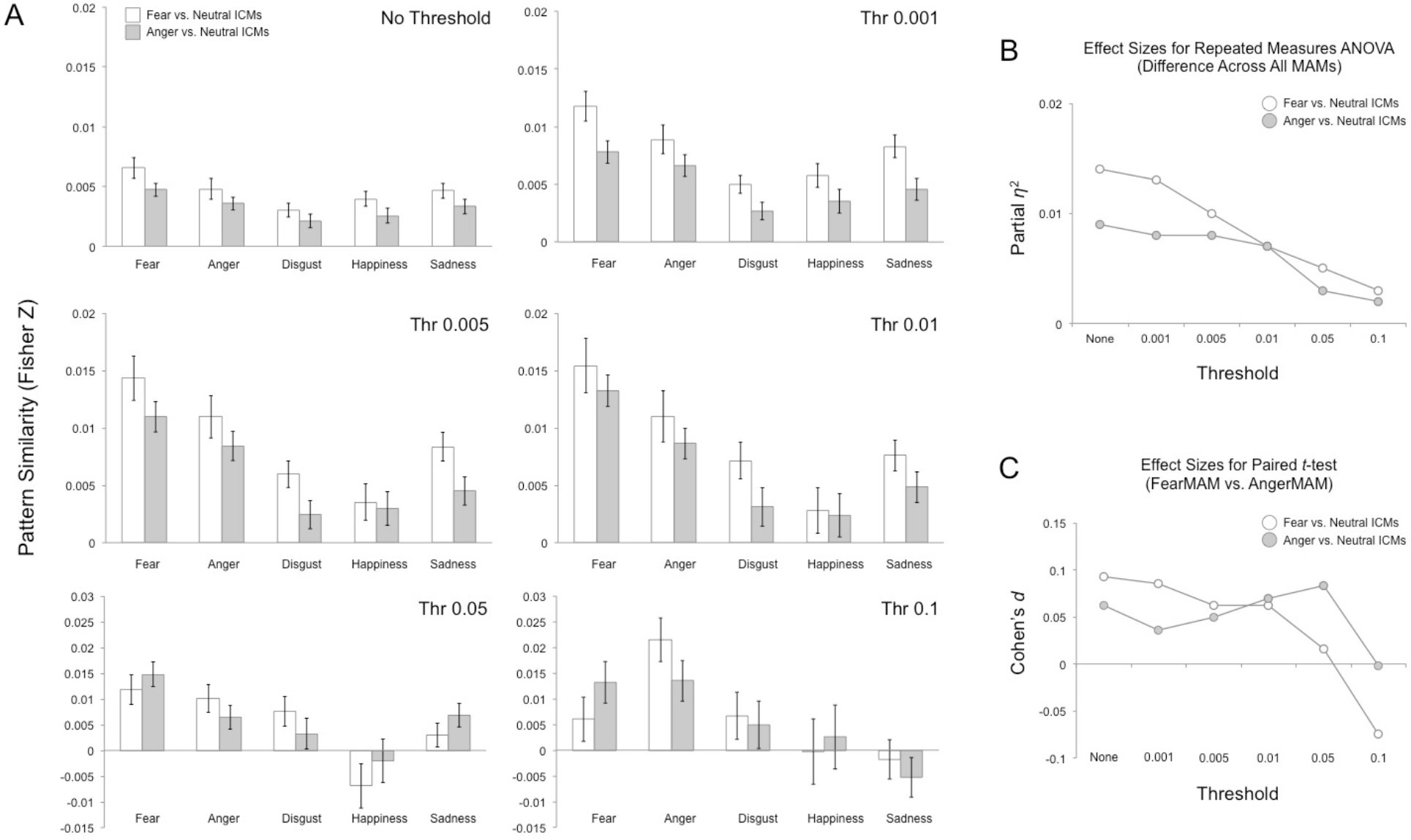
Whole brain MAM-ICM pattern similarity for fear and anger. (A) Pattern similarity measures of ICM-F (white) and ICM-A (gray) to each of the five MAMs, summarized by varying threshold levels. Overall, both fear ICMs showed greater pattern similarity with the fear MAM, but this effect disappeared when the threshold was sufficiently high (e.g., 0.1). (B) Plotting the effect sizes from repeated measures ANOVA showed a gradually declining trend as a function of increased threshold levels. (C) A similar trend was found when the effect sizes from paired *t*-tests were plotted.

### Whole Brain MAM-ICM Pattern Similarity for Anger

Again, there was significant pattern similarity between ICM-A and MAM-A (*M* = 0.0036, *SD* = 0.02, [min, max] = [−0.06, 0.06]; one-sample *t*-test: *t*_(1262)_ = 6.71, *p* < 0.000001, *d* = 0.19). This effect persisted regardless of differences in the thresholds applied to the MAMs.

Repeated measures ANOVA showed significant differences of MAM-ICM pattern similarity for ICM-A across the five MAMs (*F*_(4,5048)_ = 11.46, *p* < 0.000001; partial *η*^2^ = 0.009). *Post hoc* analysis revealed, however, that ICM-A showed greatest pattern similarity to MAM-F compared to the other four MAMs, including anger (all *p*s < 0.03). This finding remained when the threshold was increased to 0.05 (i.e., less voxels were selected); however, this effect was no longer observable when the threshold was increased to 0.1. Paired *t*-tests showed similar findings as the ANOVA, such that pattern similarity between ICM-A and MAM-A was significantly less than for ICM-A and MAM-F (*t*_(1262)_ = −2.2, *p* = 0.028, *d* = 0.06). Again, this effect was observable up until the threshold was increased to 0.05. When the threshold was set to the highest level (0.1), this effect was no longer present.

Effect sizes of the ANOVA results gradually decreased as a function of increased threshold levels. Effect sizes of the *t*-tests showed a less clear but consistent pattern where the highest threshold (0.1) yielded the smallest effect size. However, as ICM-A showed the highest pattern similarity to MAM-F and not MAM-A, diminishing effect sizes indicate a relative decrease in the specificity of ICM-A to MAM-F. These findings are summarized in Figure 4 (gray bars and circles).

### Amygdala MAM-ICM Pattern Similarity for Fear and Anger

Repeated measures ANOVA showed significant differences in pattern similarity in amygdala activity between ICM-F and all five MAMs (*F*_(4,5048)_ = 2.86, *p* = 0.022; partial *η*^2^ = 0.002). However, *post hoc* analysis showed that this effect was driven by an unexpected pattern similarity between ICM-F and the MAM for happiness. This finding remained the same when the analysis was restricted to basolateral or centromedial subregions of the amygdala. Paired *t*-tests showed similar findings as the ANOVA, such that the pattern similarity between ICM-F and MAM-F was no different from those for MAM-A. Again, this effect remained the same for basolateral or centromedial subregions of the amygdala.

Similarly, amygdala activity from ICM-A significantly differed from all five MAMs (*F*_(4,5048)_ = 4.07, *p* = 0.003; partial *η*^2^ = 0.003), but the happiness MAM showed the greatest degree of pattern similarity to ICM-A. This effect remained when only the basolateral or centromedial amygdala voxels were considered in the analysis. Once again, pairwise comparison between MAM-F and MAM-A yielded no significant differences in overall or subregional amygdala activity.

The very small effect sizes of the ANOVA and *t*-test results of the amygdala analyses showed that they were comparable to the whole brain analyses using the highest threshold levels, and not useful in parsing the distinct emotion categories. These findings are summarized in Figure 5.

**Figure 5.**
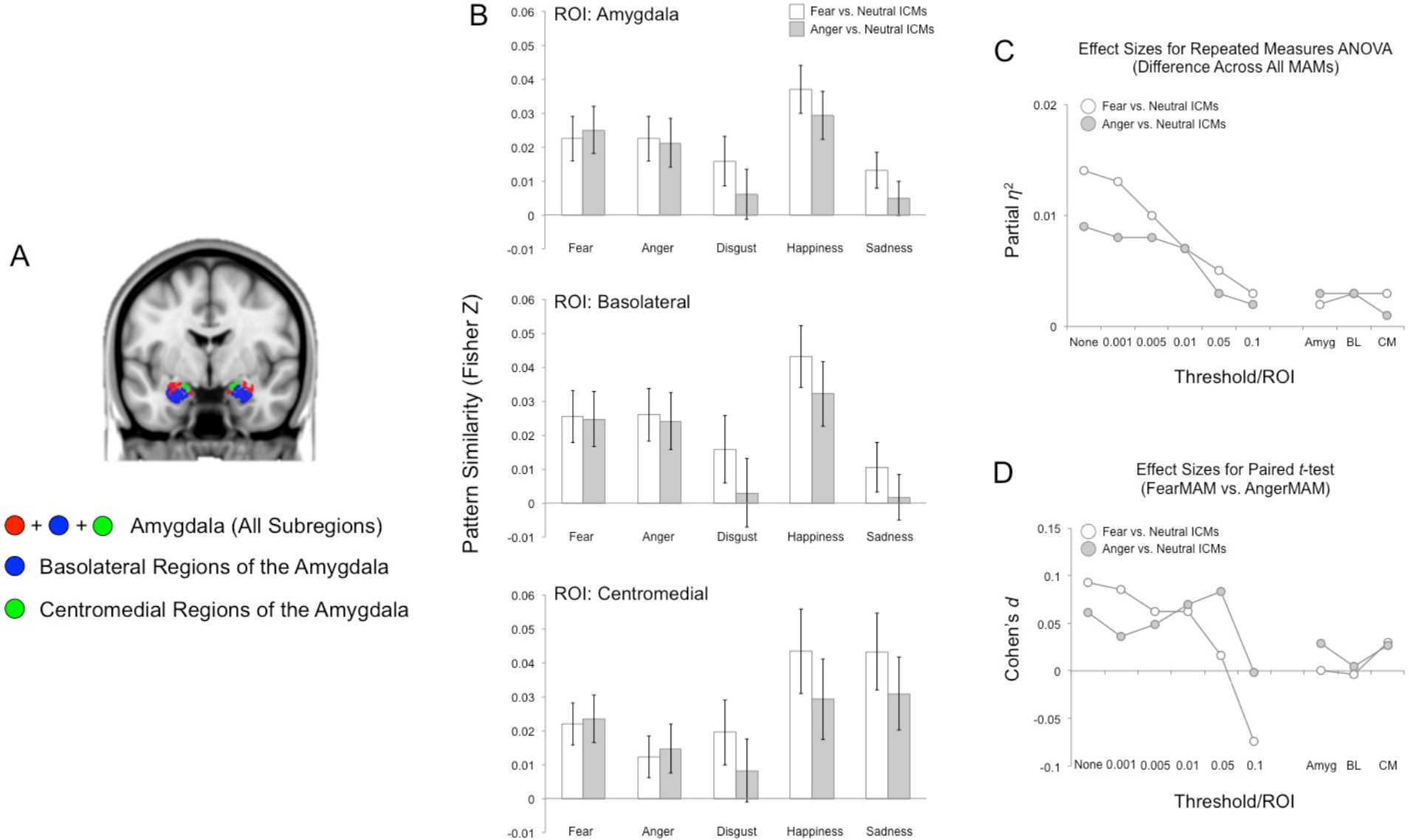
MAM-ICM pattern similarity for amygdala activity. (A) Anatomical definitions of the amygdala, basolateral amygdala, and centromedial amygdala used to select the voxels for ICM analysis (Tyszka & Pauli, 2016). (B) Pattern similarity measures for ICM-F (white) and ICM-A (gray) and each of the five MAMs, summarized by the ROIs used to define the amygdala and its subregions. Overall, neither ICM-F nor ICM-A exhibited pattern similarity with either MAM-F or MAM-A. (C) Plotting the effect sizes from repeated measures ANOVA showed a very small overall effect, regardless of activation locus (i.e., whole amygdala or subregion). The line graphs on the left represent the findings from the whole brain pattern similarity analysis (i.e., same as Figure 4B). (D) Similar findings were observed with the very small effect size for the pairwise *t*-tests.

### Non-Amygdala MAM-ICM Pattern Similarity for Fear and Anger

Nearly identical results as the whole brain MAM-ICM pattern similarity analysis were observed when the main analyses were repeated after excluding amygdala voxels (Figure 6).

**Figure 6.**
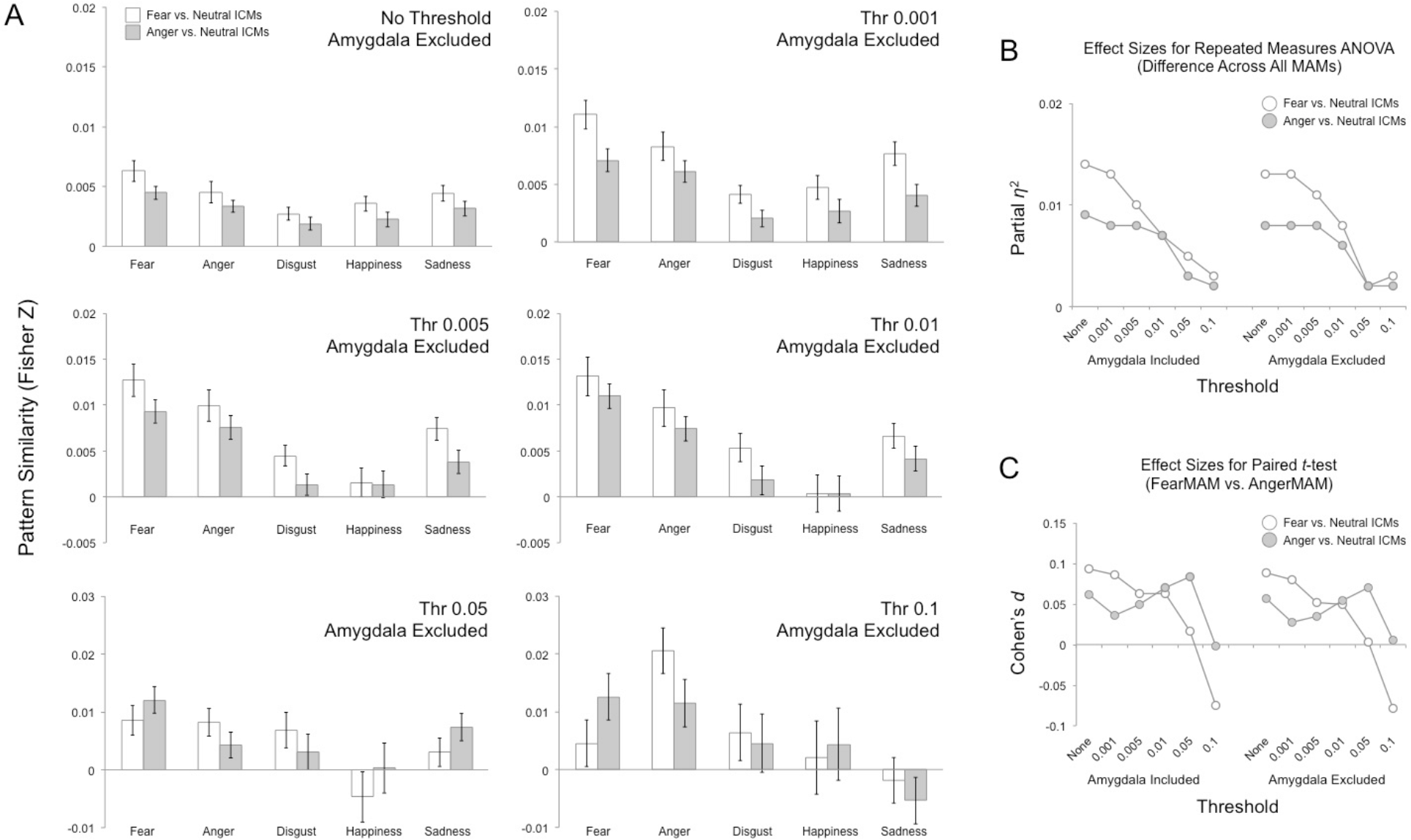
Non-amygdala MAM-ICM pattern similarity. (A) Pattern similarity measures for ICM-F (white) and ICM-A (gray) and each of the five MAMs, summarized by varying threshold levels. Overall, both ICM-F and ICM-A showed nearly identical results as the whole brain MAM-ICM pattern similarity analysis that included the amygdala. (B) Plotting the effect sizes from repeated measures ANOVA showed a gradually declining trend as a function of increased threshold levels. The line graphs on the left depict the results using whole brain voxels (i.e., amygdala included), and the line graphs on the right show the results using all non-amygdala voxels. (C) A similar trend was found when the effect sizes from paired *t*-tests were plotted.

## Discussion

Here, we compared MAMs generated for five distinct categories of emotion with fear and anger ICMs from a large study sample. Contrary to our hypothesis, we found that both ICMs exhibited the greatest pattern similarity to fear MAMs relative to all other MAMs including anger. The degree of pattern similarity decreased as the number of voxels included in the computation of the MAMs became more selective (i.e., decreased) suggesting that more distributed patterns of brain activity are better reflective of a specific emotion category. Furthermore, amygdala activity associated with either ICM was neither sufficient nor necessary for determining the overall pattern similarity between the ICMs and MAMs.

As predicted, MAM-ICM pattern similarity for fear and anger was significantly greater than zero but generally weak. This may reflect the heterogeneity inherent to the MAMs in comparison with the ICMs. The MAMs were generated from multiple studies that have used heterogenous stimuli (faces, pictures, words, films, sounds, etc.), whereas the ICMs were strictly based on facial expressions. Thus, the significant yet weak overall correlation between a given MAM-ICM pair is not surprising, as it may be partly attributable to this qualitative difference across the maps; another plausible reason is the inclusion of likely non-informative, noisy voxels present in the initial analysis with unthresholded MAM-ICM pairs. Regardless, ICM-F did exhibit the greatest pattern similarity with MAM-F as expected. This implies that ICM-F does capture a putative “fear” signature embedded within distributed brain activity, and provides support for the research strategy employed in the present study.

However, ICM-A did not exhibit the greatest pattern similarity with MAM-A. In fact, ICM-A exhibited the greatest pattern similarity to MAM-F. This suggests that the distributed brain activity associated with ICM-A is more similar to a “fear” signature than an “anger” signature. While this would appear to be paradoxical, a plausible explanation can be offered. The key here is the use of angry facial expressions with directed eye-gaze in our emotional face matching task. Angry faces with eye-gaze oriented toward the perceiver by default signal an impending aggression on part of the expressor (Adams et al., 2003; N’Diaye et al., 2009). From the perspective of the perceiver, the primary signal being communicated via anger faces is an increase in the probability of impending threat, not unlike fear faces (Whalen et al., 2001). It follows then that the perceiver’s typical response to such angry faces would better align with a threat-related response that is reminiscent of fear more so than anger. Since the MAM-A was generated from individual studies employing not just facial expressions but also other anger-*inducing* stimuli, it can be understood as representing a neural signature for anger *per se*, not the response to someone else’s anger directed at the perceiver. Thus, the present results showing that ICM-A is more similar to a “fear” than an “anger” signature could be consistent with brain activity in response to interpersonal threat (i.e., angry faces with gaze directed toward the perceiver).

These MAM-ICM pattern similarity results were dependent on the number of voxels that were included in the analysis. By systematically manipulating the number of selected voxels, we found that, in general, more voxels yielded better outcomes. However, since the inclusion of all voxels in the brain would necessarily contain those without any informational value (reflected as weak overall MAM-ICM correlations in the unthresholded analysis), additional considerations were warranted. An initial survey of the effect sizes as well as the size of the pattern similarity metrics suggested that a light threshold (0.001-0.005) provides the optimal solution, which still covers a wide range of cortical and subcortical brain regions. The distributed nature of these most informative voxels is consistent with the predictions of the original meta-analysis study (Wager et al., 2015) and generally in line with the constructionist view of emotion (Barrett & Russell, 2015; Lindquist et al., 2012), as well as findings from MVPA research on distinct emotion categories (Kassam et al., 2013; Kragel & Labar, 2015; Peelen et al., 2010; Saarimäki et al., 2016). Our data offer another piece of evidence that information about emotion categories are distributed, not localized in brain activity.

This interpretation of the present findings is furthered by the amygdala ROI analyses. If we suppose that all of the important information regarding emotion categories was being represented within the amygdala, then restricting the analysis only to the amygdala voxels should have yielded the same MAM-ICM pattern similarity outcomes from the whole brain analyses. Our data did not support this supposition, further reinforcing the main result that the inclusion of more voxels across the brain was generally beneficial in matching ICMs with MAMs. Relatedly, it is noteworthy that both MAM-F and MAM-A are characterized by similar patterns of amygdala activity (Wager et al., 2015). This suggests the possibility that this shared feature of the MAM-F and MAM-A may drive the pattern similarity with the corresponding ICMs, as both ICM-F and ICM-A are also characterized by increased amygdala activity. Our data rejected this possibility, as excluding the amygdala voxels from the analyses did not change the overall results. In fact, the findings remained remarkably similar to the whole brain MAM-ICM pattern similarity findings, with minimal changes in *z* scores and effect sizes. This illustrates that the amygdala voxels did not contribute to distinguishing discrete emotion categories in a significant way, and thus the informational value of the amygdala, at least by itself, was negligible.

The present study is not without limitations. The experimental task from which the ICMs were derived exclusively used facial expressions as the emotional stimuli. While facial expressions are widely used in the literature to examine brain responses to emotion (Costafreda et al., 2008), affective information is represented in the brain in both modality-specific and modality-general manner (Chikazoe et al., 2014; Kim et al., 2017; Shinkareva et al., 2014); thus, testing the generalizability of the present findings using ICMs derived from other modalities is warranted. It is worth again noting that the MAMs used in the present study were generated using individual studies that utilized heterogenous stimuli to represent or elicit emotions (e.g., faces, pictures, films, words). Thus, the resulting MAMs may be capturing modality-general signatures of emotions in the brain. Also, we were only able to focus on the two threat-related emotions (fear, anger), as our emotional face matching task did not include the three other emotion categories for which there are MAMs. As such, it remains to be seen whether ICMs of other emotions (disgust, happiness, sadness) would show similar mappings onto corresponding MAMs. Finally, findings from meta-analyses (i.e., MAMs) are inherently bound by the quality of the individual data (Wager et al., 2015). As technical advances in fMRI data acquisition and processing have been made in recent years, it would be worthwhile revisiting the current research topic when updated MAMs that include post-2011 studies become available.

These limitations notwithstanding, our current findings highlight that widely distributed patterns of brain activity from ICMs of threat-related emotions, across multiple brain regions and systems, may be best suited for capturing emotion categories identified by MAMs. In contrast, the amygdala was neither sufficient nor necessary for observing such MAM-ICM pattern similarity effects across discrete emotion categories. More generally, the present study offers a strategy that could further boost the utility of MAMs, whose importance has become increasingly recognized in neuroimaging research. As evidenced by the better correspondence of ICM-A to a putative signal of fear rather than anger, MAMs may be able to further shed light on the underlying mental processes captured by ICMs, which can contribute to better interpretations of findings using contrast-based task fMRI.

## Acknowledgements

We thank the Duke Neurogenetics Study participants as well as the staff of the Laboratory of NeuroGenetics. We also thank the original authors of the meta-analysis maps for their generosity in making them available for use. The Duke Neurogenetics Study was supported by Duke University and National Institutes of Health (NIH) grants R01DA031579 and R01DA033369. ARH is further supported by NIH grant R01AG049789. The Duke Brain Imaging and Analysis Center’s computing cluster, upon which all DNS analyses heavily rely, was supported by the Office of the Director, NIH under Award Number S10OD021480.

## References

Adams, R. B., Gordon, H. L., Baird, A. A., Ambaday, N., & Kleck, R. E. (2003). Effects of gaze on amygdala sensitivity to anger and fear faces. Science, 300, 1536.

Adolphs, R., Tranel, D., Damasio, H., & Damasio, A. R. (1995). Fear and the human amygdala. Journal of Neuroscience, 15, 5879–5891.

Barrett, L. F., & Russell, J. A. (2015). The Psychological Construction of Emotion. New York: Guildford.

Barrett, L. F., & Satpute, A. B. (2019). Historical pitfalls and new directions in the neuroscience of emotion. Neuroscience Letters, 693, 9–18.

Chikazoe, J., Lee, D. H., Kriegeskorte, N., & Anderson, A. K. (2014). Population coding of affect across stimuli, modalities and individuals. Nature Neuroscience, 17, 1114–1122.

Costafreda, S. G., Brammer, M. J., David, A. S., & Fu, C. H. Y. (2008). Predictors of amygdala activation during the processing of emotional stimuli: A meta-analysis of 385 PET and fMRI studies. Brain Research Reviews, 58, 57–70.

Cox R. W. (1996). AFNI: software for analysis and visualization of functional magnetic resonance neuroimages. Computational Biomedical Research, 29, 162–173.

Davis, M., & Whalen, P. J. (2001). The amygdala: vigilance and emotion. Molecular Psychiatry, 6, 13–34.

Eickoff, S. B., Laird, A. R., Grefkes, C., Wang, L. E., Zilles, K., & Fox, P. T. (2009). Coordinate-based activation likelihood estimation meta-analysis of neuroimaging data: A random-effects approach based on empirical estimates of spatial uncertainty. Human Brain Mapping, 30, 2907–2926.

Ekman, P. (1992). An argument for basic emotions. Cognition and Emotion, 6, 169–200.

Ekman, P. F., & Friesen, W. V. (1976). Pictures of facial affect. Palo Alto: Consulting Psychologists Press.

Fitzgerald, D. A., Angstadt, M., Jelsone, L. M., Nathan, P. J., & Phan, K. L. (2006). Beyond threat: amygdala reactivity across multiple expressions of facial affect. Neuroimage, 30, 1441–1448.

Greve, D. N., & Fischl, B. (2009). Accurate and robust brain image alignment using boundary-based registration. Neuroimage, 48, 63–72.

Hariri, A. R. (2009). The neurobiology of individual difference in complex behavioral traits. Annual Reviews of Neuroscience, 32, 225–247.

Kassam, K. S., Markey, A. R., Cherkassy, V. L., Loewenstein, G., & Just, M. A. (2013). Identifying emotions on the basis of neural activation. PLoS ONE, 8, e66032.

Kim, J., Shinkareva, S. V., Wedell, D. H. (2017). Representations of modality-general valence for videos and music derived from fMRI data. Neuroimage, 148, 42–54.

Kim, M. J., Loucks, R. A., Palmer, A. L., Brown, A. C., Solomon, K. M., Marchante, A. N., & Whalen, P. J. (2011). The structural and functional connectivity of the amygdala: From normal emotion to pathological anxiety. Behavioural Brain Research, 223, 403–410.

Kim, M. J., Scult, M. A., Knodt, A. R., Radtke, S. R., d’Arbeloff, T. C., Brigidi, B. D., …& Hariri, A. R. (2018). A link between childhood adversity and trait anger reflects relative activity of the amygdala and dorsolateral prefrontal cortex. Biological Psychiatry: Cognitive Neuroscience and Neuroimaging, 3, 644–649.

Klein, A., Andersson, J., Ardekani, B. A., Ashburner, J., Avants, B., Chiang, M-C., …& Parsey, R. K. (2009). Evaluation of 14 nonlinear deformation algorithms applied to human brain MRI registration. Neuroimage, 46, 786–802.

Kober, H., & Wager, T. D. (2010). Meta-analysis of neuroimaging data. Wiley Interdisciplinary Reviews: Cognitive Science, 1, 293–300.

Kragel, P. A., & LaBar, K. S. (2015). Multivariate neural biomarkers of emotional states are categorically distinct. Social Cognitive and Affective Neuroscience, 10, 1437–1448.

Kragel, P. A., Reddan, M. C., LaBar, K. S., & Wager, T. D. (2019). Emotion schemas are embedded in the human visual system. Science Advances, 5, eaaw4358.

Lindquist, K. A., Wager, T. D., Kober, H., Bliss-Moreau, E., & Barrett, L. F. (2012). The brain basis of emotion: a meta-analytic review. Behavioral and Brain Sciences, 35, 121–143.

Maren, S. (2001). Neurobiology of Pavlovian fear conditioning. Annual Reviews of Neuroscience, 24, 897–931.

N’Diyae, K., Sander, D., & Vuilleumier, P. (2009). Self-relevance processing in the human amygdala: gaze direction, facial expression, and emotion intensity. Emotion, 9, 798–806.

Nichols, T. E. (2017). Notes on creating a standardized version of DVARS. arXiv, 170401469.

Peelen, M. V., Atkinson, A. P., Vuilleumier, P. (2010). Supramodal representations of perceived emotions in the human brain. Journal of Neuroscience, 30, 10127–10134.

Phan, K. L., Wager, T., Taylor, S. F., & Liberzon, I. (2002). Functional neuroanatomy of emotion: a meta-analysis of emotion activation studies in PET and fMRI. Neuroimage, 16, 331–348.

Phelps, E. A., & LeDoux, J. E. (2005). Contributions of the amygdala to emotion processing: from animal models to human behavior. Neuron, 48, 175–187.

Power, J. D., Mitra, A., Laumann, T. O., Snyder, A. Z., Schlaggar, B. L., & Petersen, S. E. (2014). Methods to detect, characterize, and remove motion artifact in resting state fMRI. Neuroimage, 84, 320–341.

Saarimäki, H., Gotsopoulos, A., Jääskeläinen, I. P., Lampinen, J., Vuilleumier, P., Hari, R., …&, Nummenmaa, L. (2016). Discrete neural signatures of basic emotions. Cerebral Cortex, 26, 2563–2573.

Shahane, A. D., Lopez, R. B., & Denny, B. T. (2019). Implicit reappraisal as an emotional buffer: Reappraisal-related neural activity moderates the relationship between inattention and perceived stress during exposure to negative stimuli. Cognitive, Affective, & Behavioral Neuroscience, 19, 355–365.

Shinkareva, S. V., Wang, J., Kim, J., Facciani, M. J., Baucom, L. B., & Wedell, D. H. (2014). Representations of modality-specific affective processing for visual and auditory stimuli derived from fMRI data. Human Brain Mapping, 35, 3558–3568.

Tyszka, J. M., & Pauli, W. M. (2016). *In vivo* delineation of subdivisions of the human amygdaloid complex in a high‐resolution group template. Human Brain Mapping, 37, 3979–3998.

Wager, T. D., Kang, J., Johnson, T. D., Nichols, T. E., Satpute, A. B., & Barrett, L. F. (2015). A Bayesian model of category-specific emotional brain responses. PLoS Computational Biology, 11, e1004066.

Whalen, P. J., Shin, L. M., McInerney, S. C., Fischer, H., Wright, C. I., & Rauch, S. L. (2001). A functional MRI study of human amygdala responses to facial expressions of fear vs. anger. Emotion, 1, 70–83.

Yarkoni, T., Poldrack, R. A., Nichols, T. E., Van Essen, D. C., & Wager, T. D. (2011). Large-scale automated synthesis of human functional neuroimaging data. Nature Methods, 8, 665–670.

